# The Aryl Hydrocarbon Receptor senses the Henna pigment Lawsone and mediates Yin-Yang effects on skin homeostasis

**DOI:** 10.1101/459255

**Authors:** Laura Lozza, Pedro Moura-Alves, Teresa Domaszewska, Carolina Lage Crespo, Ioana Streata, Annika Kreuchwig, Marina Bechtle, Marion Klemm, Ulrike Zedler, Silviu Ungureanu Bogdan, Ute Guhlich-Bornhof, Anne-Britta Koehler, Manuela Stäber, Hans-Joachim Mollenkopf, Robert Hurwitz, Jens Furkert, Gerd Krause, January Weiner, António Jacinto, Ioana Mihai, Maria Leite-de-Moraes, Frank Siebenhaar, Marcus Maurer, Stefan H.E. Kaufmann

**Author notes:** Corresponding author: Stefan H.E. Kaufmann. Additional correspondence: Laura Lozza,; Pedro Moura Alves. these authors equally contributed to this work (Laura Lozza, Pedro Moura-Alves).

## Abstract

As a first host barrier, the skin is constantly exposed to environmental insults that perturb its integrity. Tight regulation of skin homeostasis is largely controlled by the aryl hydrocarbon receptor (AhR). Here, we demonstrate that Henna and its major pigment, the naphthoquinone Lawsone activate AhR, both *in vitro* and *in vivo*. In human keratinocytes and epidermis equivalents, Lawsone exposure enhances the production of late epidermal proteins, impacts keratinocyte differentiation and proliferation, and regulates skin inflammation. To determine the potential use of Lawsone for therapeutic application, we harnessed human, murine and zebrafish models. In skin regeneration models, Lawsone interferes with physiological tissue regeneration and inhibits wound healing. Conversely, in a human acute dermatitis model, topical application of a Lawsone-containing cream ameliorates skin irritation. Altogether, our study reveals how a widely used natural plant pigment is sensed by the host receptor AhR, and how the physiopathological context determines beneficial and detrimental outcomes.

## Introduction

The skin acts as an important first barrier of the body, which is constantly exposed to diverse environmental and mechanical insults, such as pollution, infection, injury and radiation, amongst others [1]. Additionally, the application of cosmetics and other agents can have a major impact on skin homeostasis [1]. Among the most widely used skin dyes, are the extracts of *Lawsonia inermis*, commonly known as Henna [2]. In traditional medicine, Henna has been widely used to treat bacterial and fungal infections, inflammation, cancer and various skin pathologies [3], but the underlying mechanisms remain insufficiently understood. Major side effects of Henna preparations are caused by the additive para-phenylenediamine (PPD) that has been associated with allergic contact dermatitis [4,5]. As natural product, Henna comprises a mixture of numerous compounds most of which are poorly characterized both chemically and functionally. The responsible pigment for the red colour after Henna application on skin, is the 1,4-naphthoquinone Lawsone, constituting 1-2 % of the leaves [6,7].

Recently, we unveiled that bacterial pigmented virulence factors, such as phenazines produced by *Pseudomonas aeruginosa* and the 1,4-naphthoquinone Phthiocol (Pht) from *Mycobacterium tuberculosis*, bind to and activate the Aryl Hydrocarbon Receptor (AhR), leading to AhR mediated immune defenses and detoxification of these virulence factors [8]. AhR is an evolutionarily conserved transcription factor widely expressed by almost all types of cells [9-11]. In its inactive state AhR resides in the cytoplasm in association with various chaperones. Upon activation, AhR binds to the AhR nuclear translocator (ARNT), and the resulting heterodimer induces the transcriptional regulation of multiple target genes, notably cytochrome P450 monooxygenases (*CYP1A1* and *CYP1B1*) and its own repressor, the AhR repressor (*AHRR*) [11]. Earlier studies of AhR functions focused on detoxification of xenobiotic ligands such as benzo[a]pyrene, an ingredient of tobacco smoke [12] and the highly toxic 2,3,7,8-tetrachlorodibenzo-p-dioxin (TCDD) [13]. The list of ligands is continuously expanding, encompassing endogenous molecules (*e.g.* tryptophan (Trp), kynurenine or formylindolo[3,2-b] carbazole (FICZ)), dietary compounds and bacteria-derived ligands, and others (*e.g.* Itraconazole, Lipoxin A4, Prostaglandin G2 and Quercetin) [8, 14-18]. In parallel with the increasing number of ligands, the biological functions attributed to this receptor are constantly growing rendering this receptor a ‘moving target’ of intense research [14, 19-21].

In the skin, AhR-mediated signals are critical in tissue regeneration, pathogenesis, inflammation and homeostasis [9,22,23] and AhR emerged as crucial player in the maintenance of skin integrity and immunity [9,11]. However, the outcome of AhR activation varies profoundly according to ligand properties, target cells and interactions with other signaling cascades [22-25]. Here, we aimed to better characterize the effects of Lawsone, defining its mechanisms with an emphasis on skin, the central target tissue of Henna. We demonstrate that the main pigment of Henna, Lawsone, activates the AhR-transcriptional program and modulates skin homeostasis and recovery after external insult. We show that Lawsone inhibits proliferation, and accelerates differentiation of keratinocytes. Specifically, experiments with human skin equivalents, zebrafish and mice, reveal that Lawsone modulates tissue homeostasis and tissue regeneration, thereby interfering with the physiological process of wound healing. Despite its detrimental effect on wound healing, Lawsone’s capacity to reduce proliferation and promote keratinocyte differentiation, in parallel to modulation of skin inflammation, renders it a promising candidate for therapy of hyperproliferative skin diseases.

## Results

### Henna and Lawsone activate the AhR pathway in keratinocytes

AhR triggering depends on the quality and quantity of the activators as well as the intrinsic characteristics of the cell types [11]. Due to its similarity with known AhR ligands (Fig. 1A), such as TCDD and the mycobacterial pigment Pht [8,24], we hypothesized that Lawsone, the main pigment from Henna, modulates AhR activity. *In silico* modeling studies predicted that all three molecules fit into the AhR binding pocket, albeit with different affinities (Fig. 1B and Fig. Supplement 1A). The key residues Thr289, His291, Phe295, Ser365 and Gln383 are involved in forming hydrogen bonds with each of the three ligands. Lawsone has similar interactions as Pht. After rescoring, the free binding energy was as follows: TCDD (ΔG Bind −47.568 kcal/mol), Pht (ΔG Bind −42.850 kcal/mol) and Lawsone (ΔG Bind −38.591 kcal/mol), with the lower value indicating a stronger binding in the ligand-receptor complex. Binding to AhR was confirmed in a previously established competition assay [8], where Lawsone was able to displace radioactivity labeled TCDD bound to AhR (Fig. Supplement 1B).

**Figure 1.**
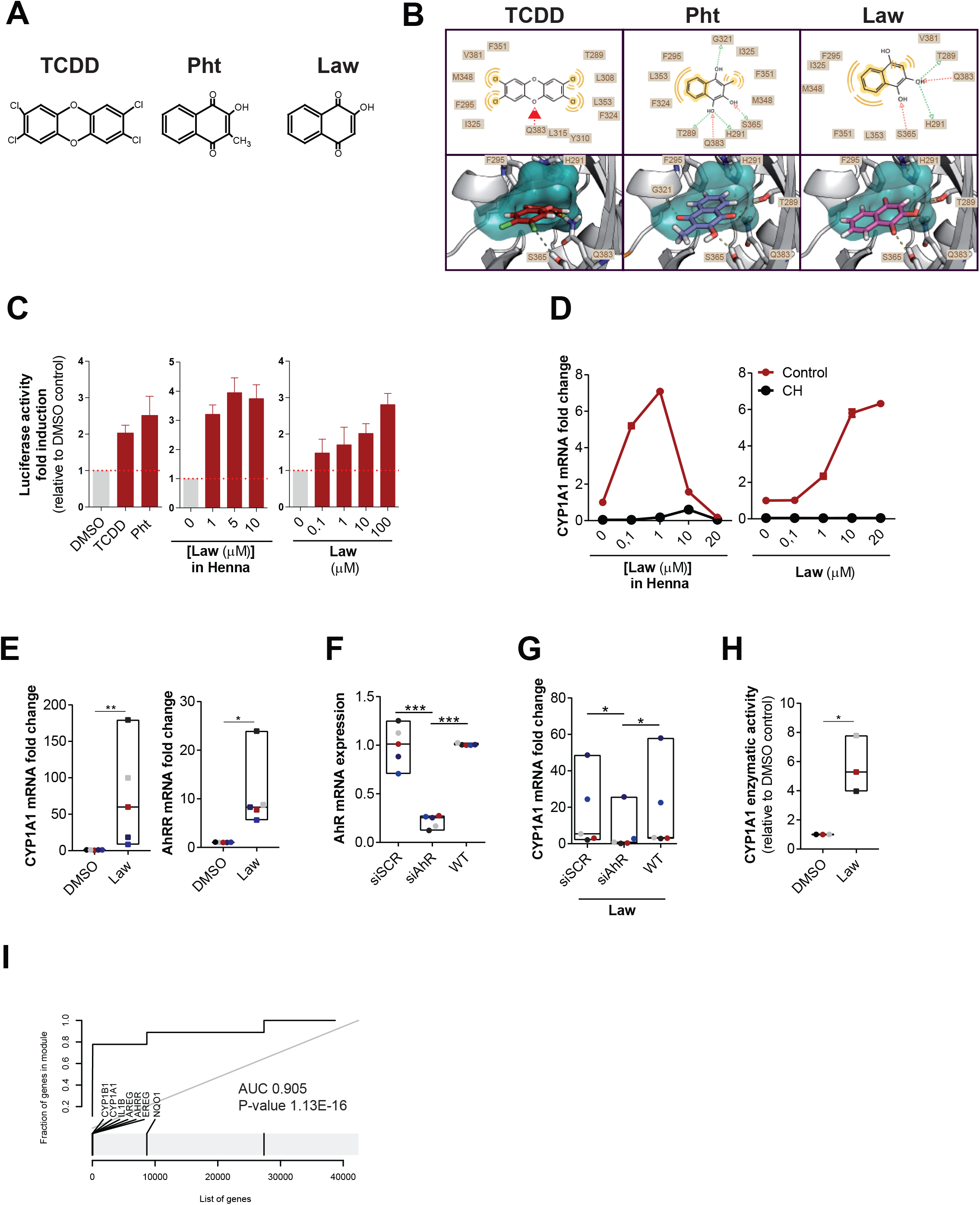
Henna and Lawsone activate AhR in HaCaT and human primary keratinocytes (**A**) Chemical structures of TCDD, Phthiocol (Pht) and Lawsone (Law) and (**B**) *in silico* modeling studies predicting binding of these molecules in the AhR binding pocket. Upper panel: 2D-interaction plot (LigandScout 4.1), hydrogen-donor (green dashed), -acceptor (red dashed), hydrophobic (orange); lower panel: 3D-interaction models, hydrogen bonds (yellow dashed), potential halogen bond (green dashed). (**C**) Luciferase activity of AhR reporter HaCaT cells stimulated for 4 hours (h) with TCDD, Phthiocol (Pht), Henna or Lawsone (Law). (**D**) Dose dependent *CYP1A1* expression of HEK cells stimulated for 4h, in the presence (black dots) or absence (red dots) of the AhR inhibitor CH223191 (CH, 12 μM) normalized to DMSO. (**E**) *CYP1A1* and *AHRR* expression after 4h Lawsone (10 μM) stimulation of HEK cells normalized to DMSO. Each dot represents one individual. (**F-G**) HEK cells were transiently transfected with AhR-siRNA (siAhR) or Scramble control (siScr) in different individuals (dots). Each color depicts results of the same individual. (**F**) AhR knockdown validation relative to non-transfected wild type (WT) cells. (**G**) *CYP1A1* expression after 4h stimulation with Lawsone normalized to DMSO. (**H**) 48 h CYP1A1 enzymatic activity in HEK cells treated with Lawsone (10 μM) normalized to DMSO. (**I**) AhR-target gene enrichment after Lawsone stimulation (10 μM) relative to TLR2 stimulation (Pam2CSK4, 300 ng/mL). Area under the curve (AUC), q-values and highly enriched genes are indicated. (**C, E-H**) Data from at least 3 independent experiments are shown. (**D**) Data from 1 representative experiment out of 2 is shown. (**C**) Mean ± S.E.M., (**D**) Mean, (**E-H**) Floating bars, Mean Min to Max. and. (**E, H**) Student’s t-test, (**F, G**) One-way ANOVA with Fisher’s test. *P<0.05; **P<0.01; ***P<0.001.

Keratinocytes are the most prominent cell type in the epidermis [23], which constitute the first contact with external agents, including Henna [7]. We developed an AhR-luciferase reporter HaCaT (immortal human keratinocyte) cell line and measured AhR activation as readout of luciferase activity after stimulation. As can be seen in Fig. 1C, both TCDD and Pht induced AhR activation in keratinocytes. Similarly, Henna and the 1,4-naphthoquinone Lawsone also activated AhR (Fig. 1C). Dose-dependent AhR activation was further confirmed in other cell types, using the AhR-luciferase reporter THP-1 (human macrophage) cell line [8] (Fig. Supplement 1C). Extending our analysis to human primary keratinocytes (HEK cells), we evaluated whether the expression of AhR target genes was differentially regulated. *CYP1A1* was induced upon stimulation with both Henna and Lawsone (Fig. 1D). AhR dependency was confirmed using the specific AhR inhibitor, CH223191 [26] (CH, Fig. 1D). *CYP1A1* transcription increased after stimulation with Henna containing 1 μM of Lawsone, while it decreased at higher concentrations (Fig. 1D, left). Henna preparations contain several components, aside from Lawsone [2], which would interfere with the kinetics of AhR activation. When cells were stimulated with Lawsone, *CYP1A1* was upregulated in a dose-dependent manner (Fig. 1D, right), without affecting cell viability (Fig. Supplement 1D-F). In keratinocytes obtained from different donors, *CYP1A1* and *AHRR* were consistently induced by Lawsone (Fig. 1E), Pht and TCDD (Fig. Supplement 1G). Notably, TLR2 stimulation (Pam2CSK4) did not activate AhR (Fig. Supplement 1G). Silencing of AhR in these cells by RNA interference (RNAi), reduced *CYP1A1* expression, further confirming that Lawsone induced *CYP1A1* in an AhR dependent manner (Figs. 1F and G and Fig. Supplement 1H). Inhibition of CYP1A1 can lead to indirect AhR activation in a mechanism involving Trp [19,27]. Using the EROD assay [28], CYP1A1 enzymatic activity was increased by Lawsone in HEK cells (Fig. 1H), as well as by the other ligands tested (Fig. Supplement 1I), thus excluding an indirect role of CYP1A1 in AhR induction in this context.

To further validate our findings, we performed microarray analysis of HEK cells stimulated with Lawsone. We identified a set of AhR dependent genes (Table 1) and visualized the gene enrichment using receiver operating characteristic (ROC) curves [29]. A high score of the area under the curve and low q value indicate a significant and specific enrichment of AhR target genes upon stimulation with Lawsone (Fig. 1I and Fig. Supplement 1J). Consistently, Ingenuity Pathway Analysis predicted the AhR canonical pathway amongst the top differentially regulated genes (Fig. Supplement 1K). Since *NQO1* can also be regulated by the transcription factor Nrf2 [30], we extended our analysis to the enrichment of genes associated with this pathway (Table 2). The area under the curve indicates that Nrf2-related genes were less enriched compared to AhR-related genes (Figs. S1L and M), pointing to a preferential activation of AhR. In summary, our results demonstrate that the 1,4-naphtoquinone Lawsone, the critical pigment in Henna, binds and activates the AhR pathway in keratinocytes. While the effects of Henna may be confounded by other components in the extract, Lawsone specifically activates AhR without causing cell toxicity, at least at the conditions tested.

**Table 1.** AhR dependent genes. The table includes AhR target genes containing the xenobiotic-responsive element (XRE) in the promoter region and genes described to be induced by AhR activation.

| Table 1. AhR dependent genes |  |  |
| --- | --- | --- |
| Name | Gene | References |
| Cytochrome P450, family 1, member A1 | CYP1A1 | Hankinson, 1995;Katiyar *et al.*, 2000;Mukhtar *et al.*, 1986 |
| Cytochrome P450, family 1, member B1 | CYP1B1 | Hankinson, 1995;Katiyar *et al.*, 2000;Mukhtar *et al.*, 1986 |
| Aryl hydrocarbon receptor repressor | AHRR | Baba *et al.*, 2001;Frericks *et al.*, 2007 |
| TCDD-inducible poly(ADP-ribose) polymerase | TIPARP | Lo and Matthews, 2012;Frericks *et al.*, 2007 |
| Interleukin-1β | IL-1β | Sutter *et al.*, 1991 |
| plasminogen activator inhibitor-2 | PAI-2 | Sutter *et al.*, 1991 |
| epiregulin | EREG | Patel *et al.*, 2006 |
| amphiregulin | AREG | Du *et al.*, 2005 |
| insulin-like growth factor 1 receptor | IGFR1 | Lo and Matthews, 2012 |
| NADP(H):quinone oxidoreductase 1 | NQO1 | Wang *et al.*, 2013 |

**Table 2.**
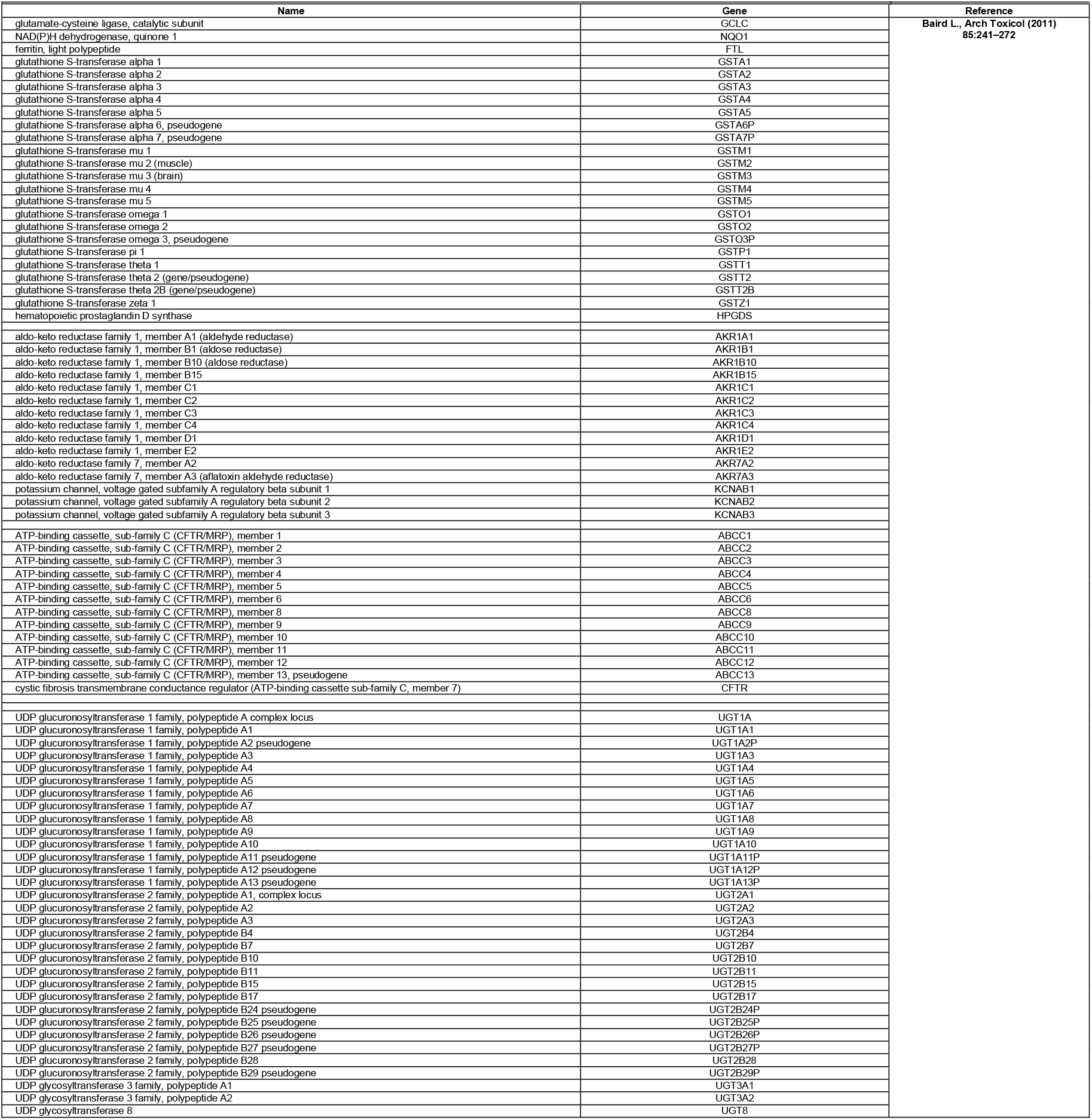
Nrf2-related genes. The table includes Nrf2 target genes.

### Lawsone stimulation modulates keratinocyte proliferation and differentiation

The AhR pathway impacts on epidermal differentiation, and the consequences of AhR activation considerably depend on the properties of the ligands and the target cells [22,25,31,32]. As demonstrated in Fig. 2A, Lawsone inhibited keratinocyte proliferation. Furthermore, microarray analysis of HEK cells stimulated with Lawsone pointed to a skewing towards differentiation (Fig. Supplement 2A). ROC curve analysis of genes of the epidermal differentiation complex (EDC), and family I and II keratins (Table 3) revealed a significant enrichment upon Lawsone stimulation (Fig. 2B). This was mainly due to upregulation of the genes involved in formation of the cornified envelope (Supplementary Dataset File 1). Cornifelin (CNFN), hornerin (HRNR), late cornified envelope 3D (LCE3D), keratin 2 (KRT2) and filaggrin 2 (FLG2) are critical for epidermal differentiation [33,34]. qRTPCR analysis confirmed the induction of these genes in HEK cells upon Lawsone exposure (Fig. 2C). Thus, Lawsone modulates the expression of genes involved in cornified envelope generation.

**Table 3.**
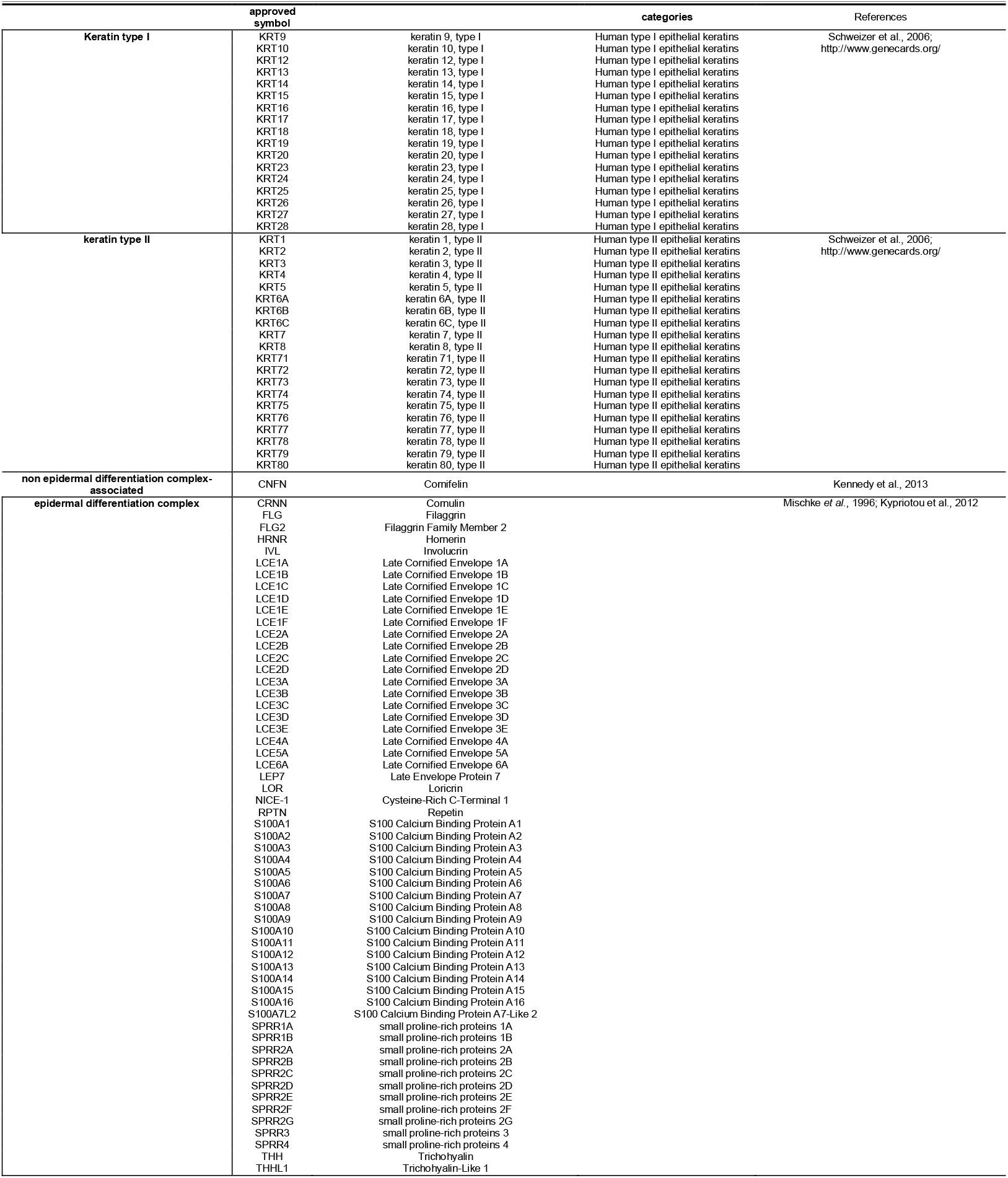
Epidermal differentiation complex and keratin genes. The table includes genes of the epidermal differentiation complex and keratins.

**Figure 2.**
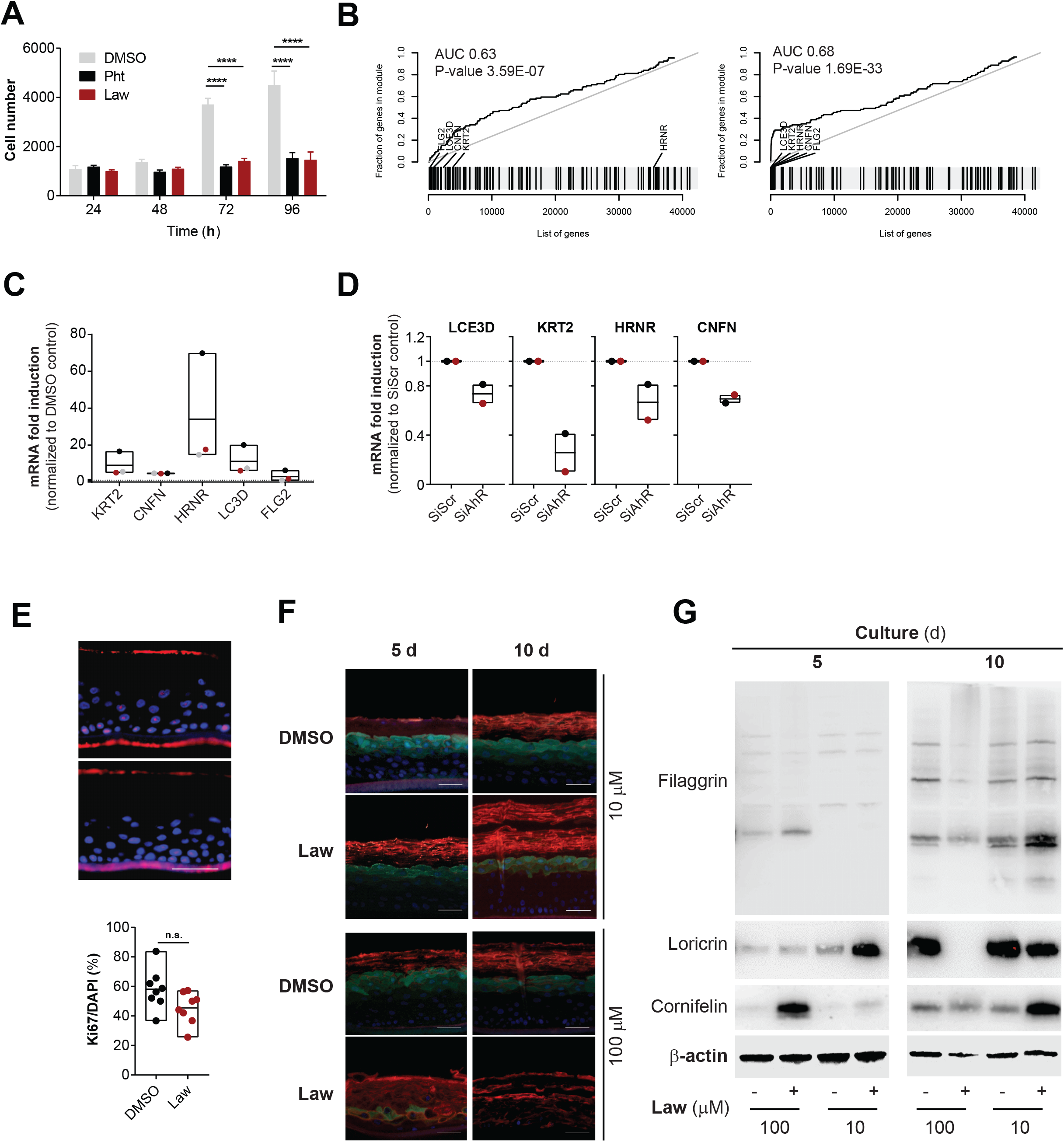
Lawsone stimulation modulates keratinocyte proliferation and differentiation (**A**) Nuc red Live 647 positive HEK cells at different time points after stimulation with Law (10 μM) and Pht (50 μM), compared to DMSO. (**B**) Epidermal differentiation complex and keratin gene enrichment of HEK cells after Lawsone stimulation (10 μM) and relative to TLR2 stimulation (Pam2CSK4, 0.236 μM) at (left) 4h and (right) 24h. Area under the curve (AUC), q-value and highly enriched genes are indicated. (**C**) *KRT2*, *CNFN*, *HRNR*, *LCE3D* and *FLG2* expression of HEK cells after 24h stimulation with Lawsone (10 μM) normalized to DMSO. Each color depicts results of the same individual. (**D**) *LCE3D*, *KRT2*, *HRNR* and *CNFN* expression on HEK cells transfected with AhR-siRNA (siAhR) or Scramble control (siScr) and further stimulated for 24h with Lawsone (10 μM). Values are relative to siScr. Each color depicts results of the same individual. (**E, top**) Epidermal skin equivalents were stimulated for 5d with Lawsone (10 μM) or DMSO and stained with DAPI (blue) and the proliferation marker KI67 (purple). (**E, bottom**) Percentage of KI67 positive cells normalized to the total number of cells (DAPI). (**F**) Representative of an *in vitro* epidermis model experiment stained for Cornifelin (red) and Loricrin (green) and (**G**) protein expression of Filaggrin, Cornifelin and Loricrin at day 5 or 10 of culture after stimulation with 10 or 100 μM of Lawsone (blots were cropped from the same gel. Full unedited gels are provided in supplementary data). (**A, C**) Data from 3 independent experiments are shown. (**D**) Data from 2 independent donors. (**E top, F, G**) One representative experiment out of 2 is shown. (**E**) Pooled data from 2 different experiments is shown. (**A**) Mean ± S.E.M., (**C, D, E bottom**) Floating bars, Mean Min to Max. (**A**) Two-way ANOVA with Fisher’s test, (**C**) One-way ANOVA with Dunn’s test. (**E, bottom**) Student’s t-test. *P<0.05; **P<0.01, ***P<0.001, ****P<0.0001.

Epidermal differentiation occurs after activation of the AP-1 transcription factor [35]. To interrogate whether epidermal differentiation requires AP-1 activity, keratinocytes were stimulated with Lawsone in the presence of the AP-1 inhibitor tanshinone IIA (TIIA) [36]. Efficient blocking of AP-1 activity was shown by inhibition of CSF3 expression (Fig. Supplement 2B) [36]. Lawsone induced upregulation of *CNFN*, *HRNR*, *LCE3D* and *KRT2* (Fig. Supplement 2C), and of the AhR-target genes *CYP1A1* and *AHRR* even in presence of TIIA indicating an AP-1 independent activation. Moreover, inhibiting AhR by RNAi reduced expression of these genes upon lawsone exposure (Figure 2D). Thus, Lawsone requires AhR activation to induce the expression of genes involved in the formation of the cornified envelope independently of AP-1 activity.

To validate our findings, we treated fresh skin biopsies from individuals after skin surgical excision with Lawsone and confirmed the upregulation of *CYP1A1* and *AHRR* (Fig. Supplement 2E), but not of *KRT2*, *CNFN*, *FLG* and *LCE3D* (Fig. Supplement 2E). We reasoned that fully differentiated skin obtained in biopsies may mask subtle differences of Lawsone on epidermal layers containing proliferating keratinocytes. Hence, we visualized epidermal differentiation over time in human epidermis equivalent models [23,34]. Keratinocytes were treated daily with Lawsone, and tissue differentiation was analyzed after 5 or 10 days of culture (Fig. Supplement 2F). As shown in Fig. 2E, the percentage of Ki67 positive cells after 5 days of treatment was slightly reduced, although not significantly, pointing to inhibition of proliferation, as observed *in vitro* (Fig. 2A). Importantly, treatment with 10 μM Lawsone increased the thickness of the *stratum corneum* after 5 and 10 days (Fig. 2F) and correlated with higher expression of loricrin (at 5 days), cornifelin (at 10 days) and filaggrin (at 10 days) measured by immunofluorescence and Western blotting (Figs. 2F and G). At higher concentrations, Lawsone further boosted the differentiation of the *stratum corneum* resulting in a disorganized epidermal structure (Fig. 2F). Hence, Lawsone impacts epidermal differentiation in human skin.

### Lawsone activates the AhR pathway in zebrafish larvae and modulates tissue regeneration

In order to further evaluate consequences of Lawsone exposure during tissue regeneration *in vivo*, we took advantage of a previously established zebrafish model [37-39]. This model organism has been extensively used in toxicology, including studies with AhR [37], as well as in skin wound healing and re-epithelization studies [38,40]. The epidermis and dermis layers occur in zebrafish larvae as early as 1 day post fertilization (dpf) [40]. 2dpf larvae were exposed to Henna Lawsone for 4 hours and AhR dependent gene expression was evaluated (Fig. Supplement 3A). Zebrafish express three isoforms of AhR (*AHR1a*, *AHR1b* and *AHR2*) [37,39] and 2 isoforms of AhRR (*AHRRa* and *AHRRb*) [40]. As in humans, the expression of *CYP1A*, as well as the repressors *AHRRa* and *AHRRb*, are regulated in an AhR dependent manner [39]. The expression of the three genes was increased upon stimulation with Henna, Lawsone (Figs. 3A and B) or TCDD (Fig. Supplement 3B). Gene induction was reversed by the AhR inhibitor, validating AhR dependency. Similar to human cells (Fig. 1H and Fig. Supplement 1I), larvae exposed to TCDD, Henna or Lawsone increased CYP1A enzymatic activity (Figs. 3C-E), which was reversed by CH223191 (Figs. 3D, E and Fig. Supplement 3C). Under these conditions, no toxicity was observed (Fig. Supplement 3D). Thus, these *in vivo* results further substantiate our *in vitro* findings demonstrating that Lawsone activates AhR signaling.

**Figure 3.**
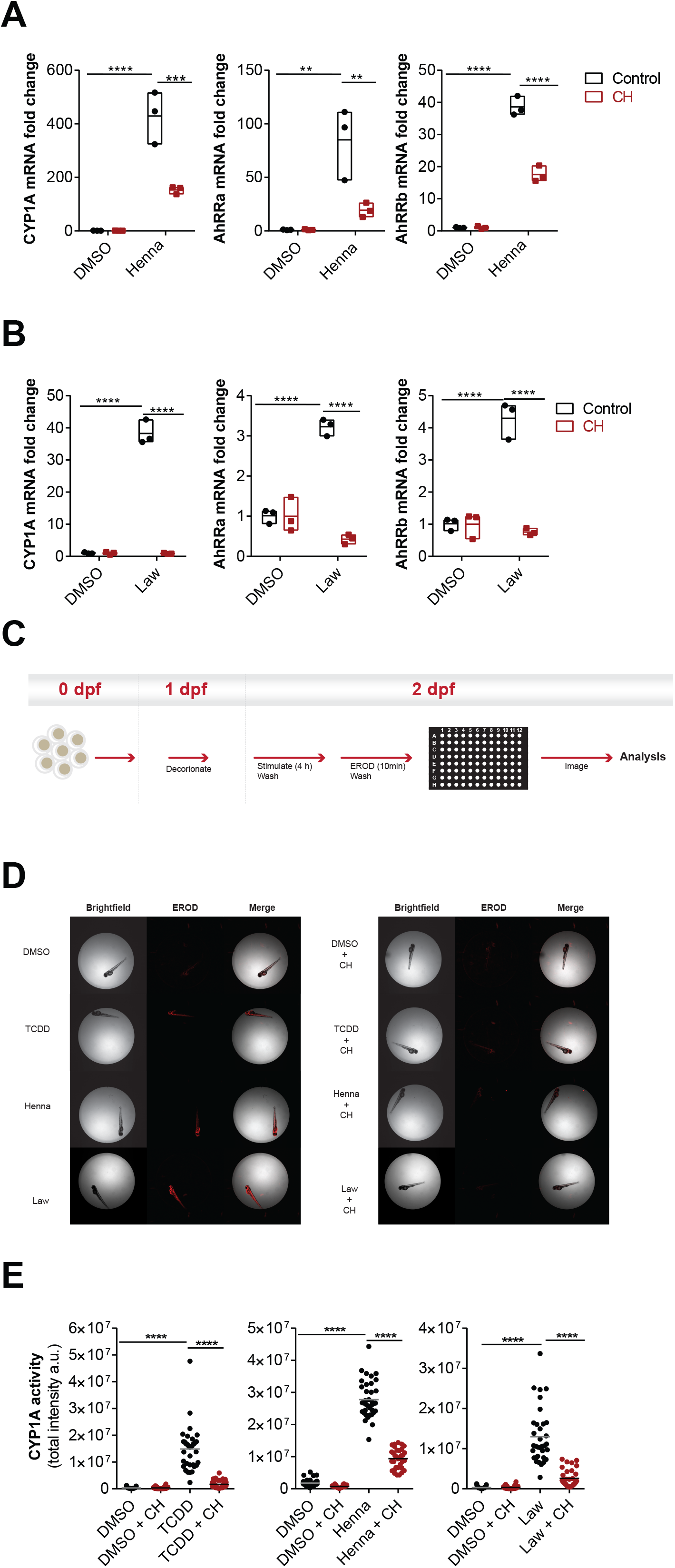
Henna and Lawsone activate AhR in zebrafish larvae (**A, B**) Fold induction of *CYP1A*, *AhRRa* and *AhRRb* transcripts from zebrafish larvae (2 days post-fertilization, dpf) treated (red squares) or not (black circles) for 2h with 5 μM of CH, followed by further 4h stimulation with (**A**) Henna (equivalent to 10 μM Lawsone), (**B**) Lawsone (10 μM) or DMSO vehicle control. Triplicates of 12 larvae depicted in each data point. (**C**) Scheme of the semi-high throughput experimental design developed to measure zebrafish larvae CYP1A enzymatic activity. (**D**) Representative images obtained upon CYP1A activity measurements using an Array Scan TM XTI Live High Content Platform. (**E**) CYP1A enzymatic activity expressed as total intensity of resorufin detected per larva (each dot represents one larva). 1 representative experiment out of 3 are shown (n=36 larvae per condition). (**A, B**) Data from 1 representative experiment out of 3 is shown. (**A, B**) Floating bars, Mean Min to Max. (**A, B**) Two-way ANOVA with Bonferroni’s test. (**E**) Two-way ANOVA with Fisher’s test. **P<0.01, ***P<0.001; ****P<0.0001.

We then performed tail fin regeneration assays and found that fin regeneration was inhibited in the presence of Lawsone (Figs. 4A and B, Fig. Supplement 4A) as observed previously with Dioxin [41,42]. Tissue damage induces the early recruitment of leukocytes to restore barrier integrity and tissue homeostasis, which critically determines the regenerative outcome [43]. Using a transgenic zebrafish line expressing GFP-labeled neutrophils (mpeg.mCherryCAAX SH378 mpx:GFP i114) [44,45] we observed that upon exposure of the tailfin wound to Lawsone, neutrophils moved (i) more randomly, (ii) for longer distances and (iii) with decreased directionality, as compared to controls (Figs. 4C-E, Fig. Supplement 4B and Movie Supplement 1). Moreover, neutrophils continued to patrol around in a “zig-zag” fashion and were not arrested at the wound (Figs. 4C, D and Movie Supplement 1). Notably, Lawsone exposure did not affect the speed of mobilizing cells (Fig. 4E). We conclude that Lawsone inhibits early steps of tissue regeneration by affecting physiological leukocyte attraction.

**Figure 4.**
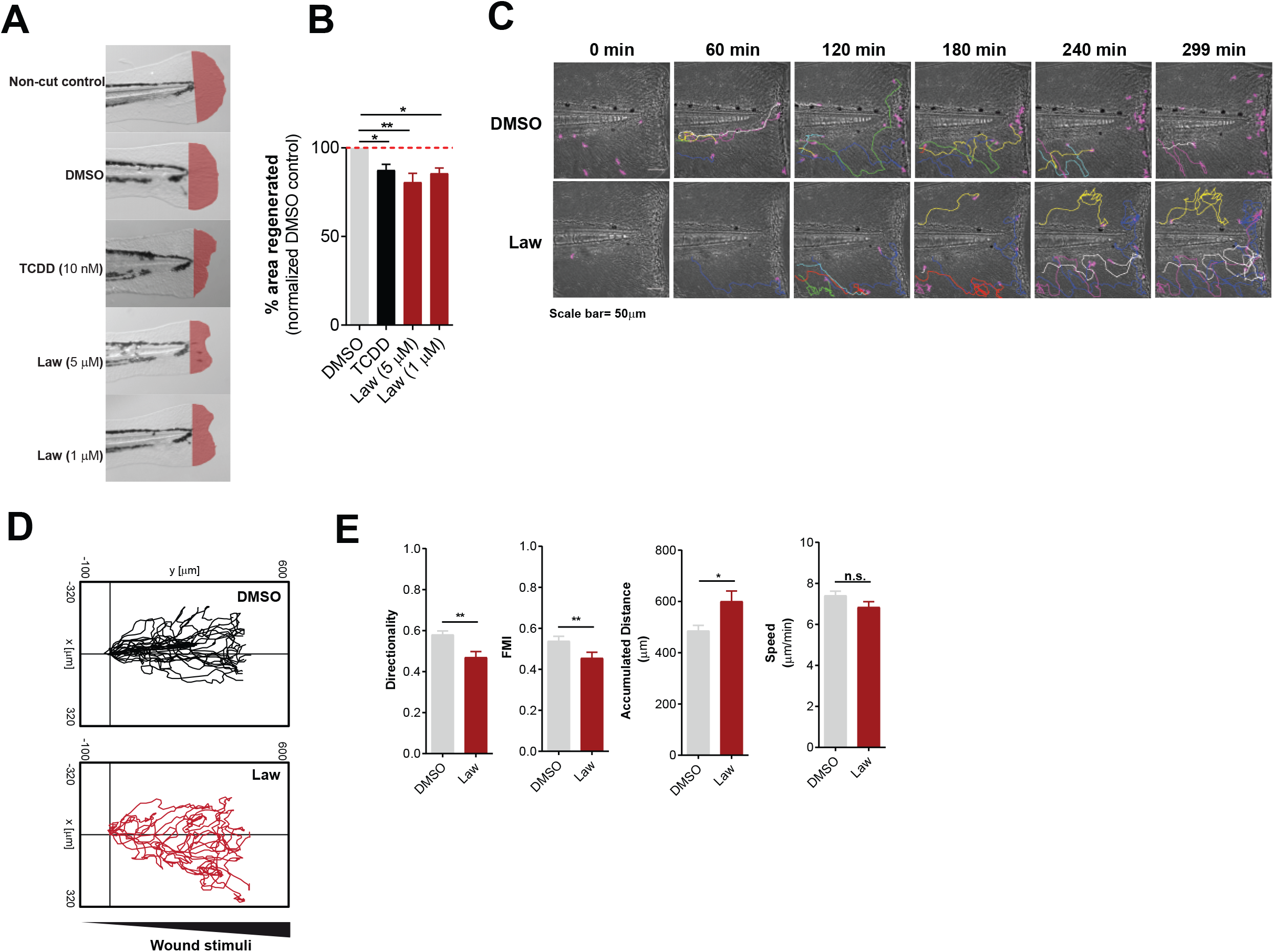
Lawsone inhibits wound healing and skin regeneration in vivo (**A**) Representative images of zebrafish fin regeneration 3 days post amputation (dpa) and exposure to different stimuli. Regenerated area depicted in red. (**B**) Quantification of the zebrafish tail fin area regenerated, normalized to DMSO treated larvae. (**C**) Neutrophil migration to zebrafish tailfin wounds visualized in DMSO or Lawsone-treated transgenic larvae Tg(mpeg.mCherryCAAX SH378 mpx:GFP i114). Frames from representative movies of migrating leukocytes in the wounded tail fin are shown. The lines indicate tracking of individual neutrophils over the indicated time point of the experiment. Wound is represented with a white dashed line. (**D**) 2D tracks of individual neutrophils migrating in the tail fin of wounded neutrophil-GFP zebrafish 3dpf larvae exposed to 10 μM Lawsone (n=8) or DMSO (n=23). (**E**) Quantification of 2D directionality, Forward migration index (FMI), accumulated distance and speed of individual leukocytes in the wounded tailfin. (**B**) Pooled data from 4 independent experiments with at least 24 larvae per condition per experiment, Mean ± S.E.M., (**E**) Data from 2 pooled experiments, Mean ± S.E.M. (**B**) One-way ANOVA with Fisher’s test, (**E**) Student’s t-test.*P<0.05; **P<0.01; ***P<0.001; n.s.-not significant.

We extended our studies to a mouse wound healing model [46]. Application of 10 μM of Lawsone on the wound for 5 consecutive days delayed wound healing (Fig. Supplement 4C and D). In sum, Lawsone interferes with the natural process of wound healing in different models.

### Lawsone ameliorates skin recovery in a model of contact skin irritation

Besides the induction of genes of epidermal differentiation, the analysis of keratinocytes stimulated with Lawsone revealed that genes related to psoriasis, dermatitis and inflammation were also affected (Table 4). Accordingly, we evaluated whether Lawsone ameliorates skin disorders characterized by irritation, inflammation and epidermal hyper-proliferation, in a human model of acute irritant contact dermatitis [47]. Skin irritation was induced by a single application of 30 μL of 5% sodium dodecyl sulfate (SDS) using self-adhesive patches which had been identified as reliable dose to induce an irritant contact dermatitis [47]. Lawsone was dissolved in base cream at different concentrations (0.5%, 1%, and 3%) and topically applied on the skin of the forearm of healthy volunteers 24h upon exposure to SDS. Images of the irritation spot and blood flux were taken daily. Decreased intensity of the flux was detected upon exposure to Lawsone, with slight differences between the concentrations and individuals tested (Fig. 5A and B). Time dependent resolution of irritation was observed in all individuals, but a strikingly faster reduction in blood flux was detected upon Lawsone exposure (Fig. 5C). Thus, Lawsone dose dependently inhibits human skin responses to irritation suggesting that detrimental or beneficial effects of Lawsone on the skin depend not only on its intrinsic nature but also on the context of skin (dys)function.

**Table 4.**
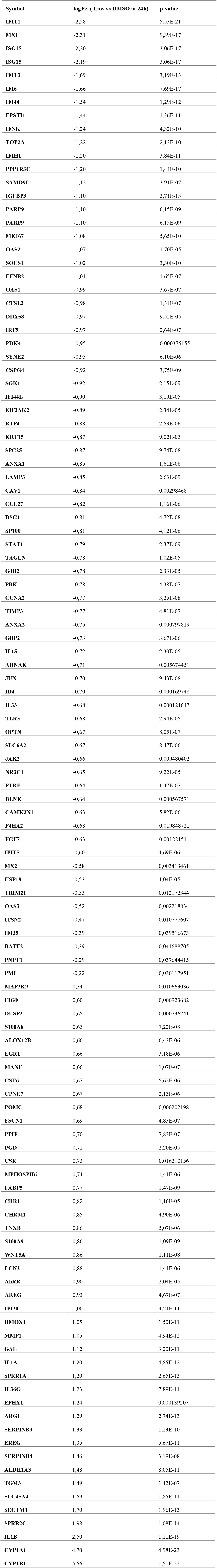
Psoriasis and dermatitis differentialy regulated genes. The table includes the genes involved in psoriasis and dermatitis that are differentialy regulated upon stimulation with Lawsone.

**Figure 5.**
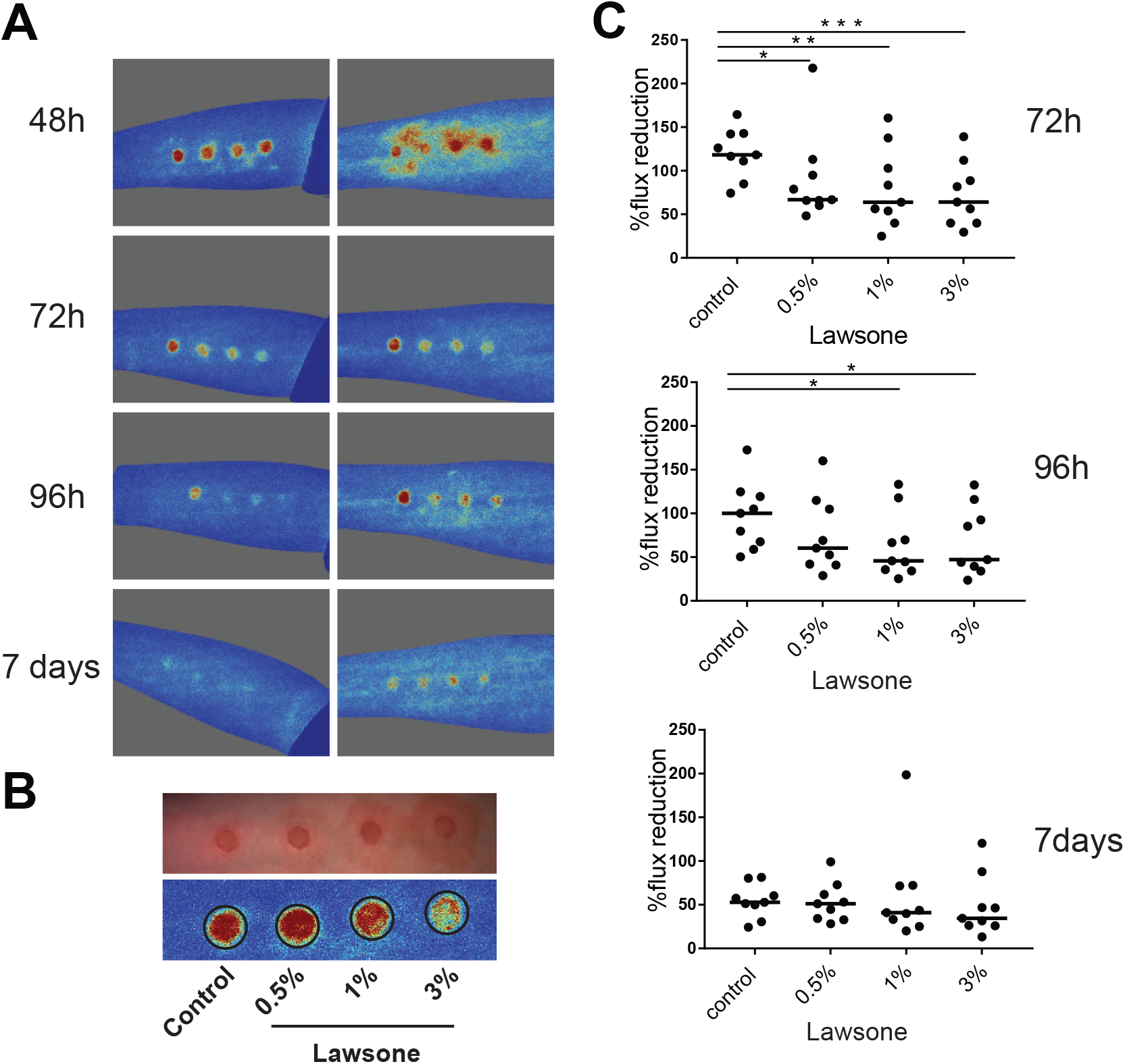
Lawsone ameliorates skin recovery in a model of contact skin irritation (**A**) Representative images of blood flux measured using the MoorFLPI-2 Full_Field Laser Perfusion Imager V1.1 software at 48-72-96 h and 7 days upon application of 0.5% SDS. Cream containing increasing concentration of Lawsone (% of Lawsone= weight of Lawsone (g) per 100g of cream) was applied 24h after SDS treatment. (**B**) Example of (top) irritation spots and (bottom) blood flux quantification. After SDS applicationall individuals were treated as follow: far left: control cream, left: 0.5 %; right 1%; far right 3% Lawsone cream. (**C**) Percentage of flux reduction at different time points normalized to the respective average flux intensity measured at 48h post-SDS application. (**A**) Representative responses of 2 out of 9 volunteers are shown. (**C**) Data from 9 individuals are shown. One-way ANOVA with Fisher’s test..*P<0.05; **P<0.01; ***P<0.001

## Discussion

Despite the widespread use of the Henna plant *Lawsonia inermis* as a cosmetic dye for hair and skin, and its broad exploitation in traditional medicine due to assumed beneficial effects, little is known about the underlying mechanisms and role of its essential pigment, Lawsone [2].

In our study, Lawsone emerged as an AhR ligand, directly binding to this receptor and eliciting AhR dependent responses in different *in vitro* and *in vivo* models. Moreover, we demonstrated that Lawsone interferes with the physiological skin regeneration processes. Lawsone modulated epidermal cell proliferation and differentiation in the skin, profoundly affecting wound healing. Nevertheless, in acute irritant contact dermatitis, Lawsone ameliorated irritation and accelerated healing.

In the skin, AhR plays a fundamental role in the maintenance of skin integrity in face of continuous environmental insults [25] and the outcome of its activation is fine-tuned by the interplay of the individual ligand properties and the physiological state of the skin [25]. Exposure to Lawsone induced the expression of AhR dependent genes not only in human primary keratinocytes and keratinocytic cell lines, but also in zebrafish larvae and human skin biopsies. AhR dependency was validated by RNAi and by using the pharmacologic AhR inhibitor CH223191. Activation of AhR can be related to inhibition of CYP1A1 activity, increasing expression of Trp metabolites activating AhR [19]. Here Lawsone did not inhibit the enzymatic activity of CYP1A, neither in zebrafish nor in human keratinocytes.

AhR has been shown to affect epidermal differentiation [22,34]. Under homeostatic conditions, AhR KO mice suffer from impaired barrier formation with enhanced transepidermal water loss and reduced expression of proteins involved in epidermal differentiation [22,32]. Similar results were obtained after exposure of keratinocytes to AhR antagonists [22], pointing to an essential role of the AhR in the physiological development of the skin barrier. Accordingly, endogenous Trp metabolites (e.g. FICZ) modulate keratinocyte functions and differentiation [15], while exogenous AhR-activators such as TCDD upregulate genes of epidermal differentiation [33,48,49]. Although FICZ and TCDD are both high-affinity AhR ligands, TCDD resists Cyp1-mediated degradation [13], while FICZ is efficiently degraded [50], suggesting that both ligand affinity and stability, shape the action on target cells. Consistent with this, TCDD favors keratinocyte differentiation but also gives rise to chloracne in overexposed humans [51], characterized by the appearance of pustules and cysts in the skin [52]. Constitutive AhR activation in keratinocytes also causes inflammatory skin lesions [53]. Hence, depending on ligand and context, AhR modulation can act as a “double-edged sword”, leading to beneficial or detrimental outcomes on skin regeneration.

In our studies, Lawsone differentially regulated distinct genes and proteins involved in keratinocyte differentiation. In agreement, proliferation of primary keratinocytes in a human organotypic skin model was decreased. Notably, the expression of specific keratinocyte differentiation genes upon Lawsone exposure was AhR dependent. Although Lawsone did not affect survival of keratinocytes, high concentrations profoundly shuffled the epidermal layers, giving rise to a thick and fragile cornified structure. Cell proliferation in regenerating zebrafish larval caudal fins in response to Dioxin has been shown to decrease [42]. Similarly, here we showed that Lawsone impairs zebrafish larval fin regeneration. Moreover, wound healing experiments in zebrafish and mouse models revealed a delay in this process caused by Lawsone. In sum, different *in vitro* (cell lines and human skin model) and *in vivo* (mouse and zebrafish) approaches conclusively demonstrate that Lawsone impacts tissue proliferation, differentiation and regeneration.

AhR mediated effects can result from different interactions between this receptor and other intracellular signaling pathways, such as Nrf2 [11,54]. Here, we showed that Lawsone upregulated the expression of the antioxidant enzyme *NQO1*, a gene also regulated by Nrf2. Nrf2 is known to protect against reactive oxygen species [30,54] and AhR and Nrf2 interactions were found crucial for the cytoprotective effects of the fungicide ketoconazole in keratinocytes [55]. In our microarray analyses, AhR dependent responses were induced more profoundly, and occurred earlier, than Nrf2 responses suggesting an important role of AhR in initiating cell responses. Yet, it is tempting to speculate that some of the elicited effects on skin may involve AhR and other molecules, such as Nrf2.

Chronic inflammatory skin disorders emerge as outcome of diverse environmental and immune factors, and diseases such as psoriasis and atopic dermatitis are characterized by dysbalanced AhR signaling. Accordingly, therapeutic interventions by AhR-targeting strategies have been suggested (25, 33, 36). For example, coal tar has been widely used for treatment of atopic dermatitis and was shown to induce AhR dependent responses in the skin [34]. Coal tar is composed of a mixture of organic compounds, and their safety and carcinogenicity have not been completely elucidated [56]. Similarly, Henna extracts contain hundreds of different components, including phenolic compounds, terpenes, steroids and alkaloids [2], but a comprehensive investigation validating the biological activities of these compounds is still missing. The effects of Henna can result from synergistic and antagonistic properties of numerous active substances. In fact, adverse events of Henna have been described, for example after ingestion and mucosal contact [57], although it appears nontoxic when applied to the skin [2]. Henna has been used for treating radiation-induced dermatitis, as well as for anti-carcinogenic, anti-microbial and anti-inflammatory purposes, although underlying mechanism and molecules involved remain elusive [2,3,58]. Given its low cell toxicity, Lawsone has clinical potential for treatment of skin disorders characterized by hyperproliferation and inflammation. Indeed, our results demonstrate that topical administration of a cream containing small amounts of Lawsone ameliorates the irritation by a chemical insult. Recently, AhR activation in keratinocytes was found to play a role in a mouse model of psoriasis by reducing inflammation [31], and current strategies to ameliorate psoriasis explore potential therapies by modulating expression of inflammatory cytokines, including IL-17 [59]. Curiously, in an Imiquimod-induced psoriasis model in mice, we observed a consistent reduction of IL-17 expression upon Lawsone topical exposure (unpublished data), pointing to potential therapeutic applications of Lawsone in skin disorders involving IL-17. Therefore, as an alternative to treatments using an undefined mixture of compounds (e.g. coal tar or Henna), we propose the Henna pigment Lawsone, and other naturally occurring naphthoquinones, as promising therapeutic candidate medicines for skin diseases. The 1,4-naphthoquinones form a family of natural pigments isolated from plants and fungi, widely used for staining food, clothing, skin and hair and in traditional medicine [60]. These include Vitamin K, Shikonin from the Chinese herb *Lithospermum erythorhizon* [61] and Juglone from the Black Walnut tree [62] that also activate AhR (unpublished data).

In conclusion, we demonstrate that the worldwide used natural product Henna and its pigment Lawsone, are sensed by AhR thereby impacting skin homeostasis. Therefore, although different AhR ligands may act as “double-edged sword” and pose harm or benefit depending on the structure and pathophysiological context, such features should be explored as future treatment options for specific dermatologic pathologies.

## Materials and Methods

### 1,4-naphthoquinone compounds and AhR agonists/antagonist

Lawsone (2-hydroxy-1,4-naphthoquinone), Dioxin (TCDD, 2,3,7,8-tetrachlorodibenzo-p-dioxin), Phthiocol (Pht, 2-hydroxy-3-methyl-1,4-naphthoquinone) were obtained from Sigma-Aldrich, and CH223191 from Santa Cruz Biotech. All compounds were solubilized in DMSO. Henna was acquired in a conventional shop and dissolved in water. To ensure that the concentration of Lawsone in the Henna preparation was comparable to that of the purified pigment employed in our experiments, we quantified the amount of Lawsone contained in the commercial Henna powder preparation by thin-layer chromatography (TLC).

### In silico homology modeling

A BLAST search with the sequence of hAhR PASB as a template revealed 58 hits in the Protein Data Bank (PDB) of experimental crystal structures. Based on sequence alignment, similarities, as well as bound ligands, 7 crystal structures were selected for a multiple sequence alignment and used to build a multiple template-based homology model of hAhR PASB. Apart from X-ray complex of HIF2α/ARNT, previously used as single template [63-65], we additionally downloaded HIF2α complexed with agonists and antagonists (PDB ID: 3F1O, 4GHI, 4GS9, 4H6J, 5TBM (chainA)), homologous complexes of HIF1α (4ZPR (chain B)) and of Clock/BMAL1 (4F3L) from the PDB and isolated the respective chains. Modeller 9.17 was used to create the multiple template-based homology model of hAhR. The resulting models were ranked by DOPE scoring. The best scoring model was selected for all subsequent modeling activities. Subsequently, model quality was checked, and the Protein Preparation wizard included in Maestro11v0 software (Schrödinger, LLC, New York, NY, 2018) was used to adjust structural defects using default values. All ligands were downloaded from Pubchem and thereafter analyzed by the Ligand Preparation Wizard to correct improper connectivities.

### In silico docking studies

Molecular docking was performed using Glide included in Maestro 11v0 software. Glide docking methodologies use hierarchical filters searching for possible ligand positions in the receptor binding-site region. Initially we set up the receptor grid defining the shape and properties of the receptor binding site important for scoring the ligand poses in later steps. Ligand flexibility was accounted by exhaustive sampling of ligand torsions during the docking process and suitable poses selected for further refinement of torsional space in the field of the receptor. Finally, in a post-docking minimization the selected poses were minimized with full ligand flexibility. The docking results were ranked by GlideScore.

The receptor grid for the hAhR homology models was set up using default parameters. Flexible ligand docking was carried out in a standard precision (SP) approach. The resulting GlideScore is an estimate of the binding affinity. Molecular mechanics application Prime MM-GBSA was used for rescoring the docking poses. MM-GBSA binding energies (MMGBSA ΔG Bind) are approximate free binding energies of protein-ligand complexes, with a more negative value indicating stronger binding.

### AhR binding studies

AhR binding experiments were performed as described previously [66]. Briefly, livers from WT mice were collected and minced in MDEG buffer (25 mM MOPS, 1 mM DTT, 1 mM EDTA and 10 % Glycerol, pH 7.5). Lysates were further homogenized, ultracentrifuged (100,000 g, 1h) and the cytosolic fraction collected. Protein concentration was determined and diluted to a final concentration of 5 mg of cytosol protein/mL. Binding studies were performed upon overnight 4°C incubation with [3H] TCDD, in the presence or absence of an excess of unlabelled TCDD. After incubation, charcoal Norit A suspension was added into the reaction mixture and incubated on ice. After centrifugation (25,000 g, 15min at 4°C), radioactivity was measured in a scintillation counter.

### Cell culture and stimulation

Human epidermal keratinocytes (HEK) (Life Technologies) were grown in Epilife medium containing human keratinocyte growth supplement (Life technologies) and 1% (v/v) penicillin– streptomycin-gentamycin (GIBCO). Cells were used between 50-70% of confluence to avoid spontaneous differentiation due to dense cultures and up to three passages. Cells were trypsinized 15 minutes (min) at 37 °C, washed with blocking buffer (PBS+ 1% FCS) at 180 g for 7 min, counted and plated overnight. HEK cells were then incubated with Lawsone or positive controls as indicated in the text, in the absence of epidermal growth factor, and analyzed at different time points. For AhR inhibition 12 μM of the AhR inhibitor CH223191 was added to HEK cells 1 hour (h) before stimulation with the AhR activators. Alternatively, HEK cells were treated for 24h with ON-TARGET plus siRNA AHR (NM_001621) and ON-TARGETplus Non-targeting Pool (Table S1, Dharmacon), according to manufacturer’s instructions. Cells were then stimulated with ligands, and *CYP1A1* transcripts analyzed after 4 or 24h. *CYP1A1* expression was normalized to glyceraldehyde-3-phosphate dehydrogenase (*GAPDH*) and results shown as fold induction (2^-δδCt^) against non-transfected cells treated with the vehicle control (DMSO).

In some experiments, cells were pretreated for 15 min with 1 μM of the the AP-1 inhibitor TIIA [36] (Sigma-Aldrich) before stimulation with AhR activators. The time was selected by measuring the inhibition of *CSF3* expression (target of AP1) [36].

HaCaT cells (Human keratinocyte cell line provided by DKFZ, Heidelberg and CLS) [67] and THP1 cells (human monocytes, ATCCTIB-202, Wesel, Germany) were grown in DMEM and RPMI 164, respectively. Both media were supplemented with 10% fetal calf serum (FCS), 1% (v/v) penicillin–streptomycin, 1% (v/v) gentamycin, 1% (v/v) sodium pyruvate, 1% (v/v) L-glutamine, 1% (v/v) non-essential amino acids, 1% (v/v) HEPES buffer and 0.05 % M2-mercaptoethanol (all reagents provided by GIBCO). Cells were kept at 37°C in 5% CO_2_. THP-1 cells were differentiated into macrophages by treatment with 200nM of phorbol-12-myristate-13-acetate (PMA, Sigma-Aldrich).

### Lentiviral infection and reporter cell line development

The construct for generation of the AhR reporter cell lines was obtained from SABiosciences (http://www.sabiosciences.com/reporter_assay_product/HTML/CLS-9045L.html). Briefly, the Cignal™ Lenti XRE Reporter is a replication incompetent, VSV-g pseudotype lentivirus expressing the firefly luciferase gene under the control of a minimal (m) CMV promoter and tandem repeats of the dioxin-responsive element (DRE). Upon stimulation of the AhR pathway, induction of luciferase expression can be used as readout of activation. Lentiviral infection was performed according to the protocols available at RNAi Consortium website (https://www.broadinstitute.org/genome_bio/trc/publicProtocols.html). 2.2×10^4^ cells per well in a 96 well plate (NUNC) were plated overnight. Following day, medium was removed and lentiviruses were added to the cells in medium containing 8 mg/ml of polybrene (Sigma-Aldrich). Plates were spun down for 90 min at 2200 rpm at 37°C. Transduced cells were further selected using puromycin (Calbiochem; 5 mg/ml) 2 days(d) after infection.

### Luciferase assay

AhR reporter cell lines were stimulated for specified time and concentration of the ligand. Cells were harvested in reporter lysis buffer (Promega) and supernatant used to determine luciferase activity using Dual-Glo Luciferase Assay System (Promega) according to manufacturer’s instructions. Luciferase activity was normalized to the amount of protein determined by Bradford reaction (Protein Assay Kit, Pierce). Results are shown as fold induction by normalizing the activation of the different compounds against non-stimulated or vehicle control.

### Ex vivo stimulation of skin biopsies

Skin was cut in small pieces (1 cm^2^) and treated 24h with vehicle control (DMSO) or 10 μM of Lawsone followed by cell disruption and lysis in Trizol.

### Stimulation and development of human epidermal skin equivalents

Undifferentiated human epidermal skin equivalents (EpiDerm model, EPI-201, MatTek Corporation) were cultured at the air–liquid interface for 10d. Cells were daily treated with 10 μM or 100 μM of Lawsone or DMSO.

### Immunostaining of HEK cells or human skin equivalents and image analysis

Cytotoxicity of Lawsone was measured by phosphorylation at Ser139 residues of the H2A.X histone. Phosphorylation of the H2A.X histone occurs at the site of DNA damage after exposure to polyaromatic hydrocarbons, hydroxyl radicals or ionizing radiation [68]. 4h after Lawsone exposure, HEK cells were fixed with 2% paraformaldeyde for 20 min at room temperature (RT) and permeabilized with 0.1% Triton for 5 min at RT. After 30 minutes in blocking buffer, cells were stained with α-phospho-Histone H2A.X (Millipore) for 1h at RT, followed by staining with the α-rabbit IgG AlexaFluor488 (Dianova,) for 1h at RT. Nuclei were stained using Nuc red Live 647 (Life technologies). Cell image acquisition and analysis was perfomed using Arrayscan XTI Live High Content Platform (ThermoFisher Scientific).

Formalin-fixed paraffin-embedded skin equivalents were stained either with hematoxylin and eosin (Sigma-Aldrich) or anti-human cornifelin (Sigma) and loricrin (Abcam), followed by anti-rabbit AlexaFluor555 and 488 respectively. Nuclei were counterstained with DAPI. Images were acquired using a Leica DMRB fluorescent microscope and analyzed with ImageJ (https://imagej.nih.gov/ij/).

### RT–PCR and RT-PCR multiplex gene expression profiling

Total RNA was extracted using 500 μL of trizol (Life technologies), followed by chloroform (1:5) and isopropanol (1:2) phase separation. RNA was washed with ethanol and resuspended in RNase free water. RNA quality and concentration were determined by spectrophotometry (Nanodrop 2000c, ThermoFischer Scientific). Complementary DNA (cDNA) synthesis was generated using Superscript III Reverse Transcriptase (Invitrogen), according to manufacturer’s instructions. Quantitative RT–PCR was performed using TaqMan master mix (Life technologies). In some experiments multiplex gene expression profiling was performed using the Biomark HD of Fluidigm as previously described [69]. Gene expression was normalized to *GAPDH*. The average threshold cycle of triplicate reactions was employed for all subsequent calculations as 2^-δδC^t relative to vehicle control (DMSO). Taqman probes (Life technologies) are listed in Table S1.

### Western blotting analysis

Proteins of human skin equivalents were isolated with radioimmunoprecipitation assay buffer and protein concentrations analyzed with the Pierce BCA Protein Assay Kit (Termo Fisher Scientific), according to the manufacturer’s instructions. Anti-human cornifelin and filaggrin were purchased from Sigma and anti-human loricrin from Abcam.

### Lactate dehydrogenase (LDH) assay

LDH was purchased from Pierce^TM^ (Thermo Scientific) and used according to the manufacturer’s instruction. Percentage (%) of cytotoxicity was calculated as:

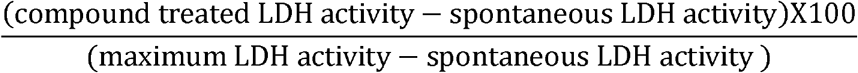

### Ethoxyresorufin-O-deethylase (EROD) activity

The enzymatic activity of CYP1A1 was used as readout of AhR activation. The EROD assay detects the CYP1A1 enzymatic activity by measuring the conversion of ethoxyresorufin into resorufin [70] in the medium of HEK cells. Briefly after 48h of stimulation with AhR activators, 4 μM resorufin ethyl ether (EROD, Sigma-Aldrich) and 10 μM dicoumarol (Sigma-Aldrich) were added to HEK culture for 1h and activity measured with the Fluoroskan Ascent Microplate Fluorometer (Thermo Labsystem). The activity was corrected to the amount of protein measured by Bradford assay and normalized to the vehicle control (DMSO).

### In vivo zebrafish experiments

Fertilized embryos were used for all experiments. One day post fertilization (dpf) larvae were manually dechorionated under a Leica MZ6 Stereomicroscope. Each experimental group consisted of 12 larvae unless stated otherwise.

#### • Larval exposure experiments

In larval exposure experiments, 2dpf AB strain larvae were exposed to different ligands for 4h, in the presence or absence of CH223191 (5 μM). After exposure, larvae were euthanized with Tricaine (MS-222, 300 μg/mL SIGMA) and placed in Trizol for RNA isolation or used for EROD experiments performed as described previously [72]. Briefly, After exposure, zebrafish larvae were washed and placed in medium containing 0.4 μg/mL of 7-ethoxyresorufin (Cayman Chemical) for 5 min. Non-fluorescent 7-ethoxyresorufin diffuses into the embryo and is O-deethylated into resorufin, a fluorescent product that can be measured [72]. Embryos were anesthetized with Tricaine (MS-222 168 μg/mL, SIGMA) [73], placed in black 96 well plates with clear bottom (Thermo Fisher) and imaged in an Array Scan ^TM^ XTI Live High Content Platform (Thermo Fisher). Brightfield images were used to identify shape of fish and fluorescence (filters excitation: 549/15 nm, emission: 590-624 nm) was determined per fish as a readout of CYP1A activation. Syber-green primers (Eurofins) are listed in Table S1

#### • Larval tail fin regeneration

2dpf AB larvae were anesthetized with Tricaine (MS-222, 200 μg/mL, Sigma) and tail fin was amputated as described previously [41]. After amputation, larvae were exposed to different ligands for 1h. After exposure, and several washes with embryo medium, larvae were kept for 3d in an incubator at 28°C with cycles of 14h of light and 10h of darkness. Afterwards, larvae were anesthetized with Tricaine (MS-222, 168 μg/mL, Sigma) and visualized in an M205 Leica stereomicroscope. Data analysis was performed on ImageJ software (https://imagej.nih.gov/ij/).

#### • Zebrafish cell migration

The transgenic line used in the study was mpeg.mCherryCAAX SH378 mpx:GFP i114, where neutrophils stably express GFP [44,45]. Imaging was performed on 3dpf larvae treated, wounded and mounted as reported previously [74]. Briefly, embryos were pretreated with 10 μM Lawsone or DMSO, in E3-tricaine solution (E3/T; Sigma; 200 μg/mL) for 1h. Fish were anaesthetized in Lawsone-containing E3/T, and a section of the tail was cut using a razor blade. Fish were then embedded lateral side down in 1% low melting point agarose (dissolved in E3/T), over MatTek glass bottom culture dishes and overlaid with the drug in E3/T. Time-lapse fluorescence images were acquired with an Andor Revolution spinning-disk confocal unit equipped with an inverted Nikon Eclipse Ti microscope and an XYZ motorized stage, coupled to an EMCCD camera (Andor) and a Yokogawa CSU-X1 scanning head and driven by Andor iQ 2.5.1 software. GFP imaging was performed using 488-nm laser line.

Image sequences were generated every minute using a 20X NA 0.75/20X Super Fluor objective and 3.44 μm step size. Bright field images were taken at low-level illumination with a halogen lamp. Where indicated, images were processed with Manual Tracking module (ImageJ software, NIH) on maximum intensity projection. Upon background subtraction for each fluorescence channel, a Gaussian blur filter was applied. Brightness and contrast were set and then multi-channel image sequences were overlaid.

#### • Zebrafish cell dynamics analysis

Neutrophils were tracked with the Manual Tracking plugin (ImageJ). The resulting 2D coordinates were analyzed using the Chemotaxis Tool plugin (Ibidi, Germany). (http://ibidi.com/fileadmin/products/software/chemotaxis_tool/IN_XXXXX_CT_Tool_2_0.pdf). Directionality of the path represents a measurement of the straightness of cell trajectories and is calculated as:

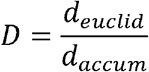

Where d_accum_ is the accumulated distance of the cell path and the d_euclid_ is the length of the straight line between cell start and end point [75].

The forward migration index represents the efficiency of the forward migration of cells towards the wound, and is calculated as:

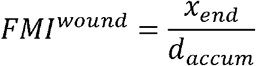

Where x_end_ is the cell end position in the axis towards the wound.

### Mouse wound healing experiments

C57BL/6 mice were bred and housed in community cages at the Animal Care Facilities of the MPIIB, Marienfelde, Berlin. Mice were used at 7-8 weeks of age. An excision of 6mm was performed at the back of the mice anesthetized with Isofluran. The wound was immediately treated with Lawsone (10 μM) or DMSO in PBS. Treatment was followed up for 5 consecutive days. Pictures were taken daily until day 6 using a Fujifilm FinePix S5800 camera. Analysis of the data was performed using ImageJ software (https://imagej.nih.gov/ij/). To calculate the size of the wounds the circumference was normalized to the length of an internal control (1cm of the ruler in the picture) and results were further normalized to day 0.

### Contact skin irritation model

Skin irritation was induced with 5% sodium dodecyl sulfate (SDS) in conserved water DAC (NRF S.6) on 4 spots of the volar forearm of 9 subjects using round self-adhesive patches with a diameter of 1.2 cm (Curatest^®^F, Lohmann & Rauscher, Germany). Patches were removed after 24 h and the skin was carefully cleaned with water. Lawsone was applied to the SDS treated sites at concentrations of 0.5%, 1% and 3% (g/g) in base creme. Pure base creme served as an intraindividual control. Treatment sites were covered by self-adhesive patches for another 24 h. The extent of skin irritation was assessed by using Moor Full-field Laser Perfusion Imager (FLPI-2, Moor Instruments, Axminster, UK) at time points 2, 3 and 7 days after induction of skin irritation.

### Microarray hybridization protocol, data preprocessing and analysis

Gene expression microarray studies were carried out as dual-color hybridization of HEK cells from one donor. RNA labeling was performed with the Quick Amp Labeling Kit, two-color (Agilent Technologies). In brief, mRNA was reverse transcribed and amplified using an oligo-dT-T7 promoter primer, T7 RNA Polymerase and Cyanine 3-CTP or Cyanine 5-CTP. After precipitation, purification, and quantification, 300 ng cRNA of both samples were pooled, fragmented and hybridized to custom-commercial whole genome human 8×60k multipack microarrays (Agilent-048908) according to the supplier’s protocol (Agilent Technologies). Scanning of microarrays was performed with 3 μm resolution using a G2565CA high-resolution laser microarray scanner (Agilent Technologies). Microarray image data were processed with the Image Analysis/Feature Extraction software G2567AA v. A.11.5.1.1 (Agilent Technologies) using default settings and the GE2_1105_Oct12 extraction protocol.

The extracted single-color raw data txt files were analyzed using R and the associated BioConductor *limma* R package [76,77] for differential expression analysis. The data set was background corrected and normalized using *loess* method. Microarray data were deposited in the NCBI’s Gene Expression Omnibus (GEO, accession number GSE99901).

We used the lmFit function to fit a linear model which included the factors stimulus type (Lawsone and Pam2CSK4) and treatment (stimulated/control) as well as an interaction term. The p-values were calculated based on moderated t statistics and most differentially regulated genes were retrieved with *topTable* function from *limma* package.

Genes associated with AhR and Nrf2 activation or keratinocyte differentiation were manually chosen on the basis of literature, and three custom gene lists were created: AhR dependent-genes (Table 1), Nrf2-related genes (Table 2) and EDC-keratin genes (Table 3). Gene set enrichment analysis was performed and visualized using R-package *tmod* for analysis of transcriptional modules [78]. In the first step, CERNO statistical test was applied to the list of genes contained in the linear fit model with *tmodLimmaTest* function. Next, ROC curve was plotted for the respective modules using *evidencePlot* function from *tmod* package [29,77] Genes presenting highest influence on the module enrichment were identified and labeled on the ROC curve. Statistical script in R including all steps of the microarray analysis can be obtained by request. Ingenuity Pathway Analysis (https://www.qiagenbioinformatics.com/products/ingenuity-pathway-analysis/, version 33559992) was performed to identify the top canonical pathways differentially regulated upon 4h stimulation of HEK cells with Lawsone (10 μM) when compared to DMSO. Pathway analysis was performed using log2 fold changes and p-values obtained from comparisons between the different stimuli.

### Statistical analysis

Statistical analysis was performed with GraphPad Prism v7.03 (GraphPad software Inc., USA). P-values were calculated using student’s t-test, One-Way or Two-Way ANOVA as stated for each experiment. The confidence interval used is 95%. P-value (P) *<0.05; **<0.01; ***<0.001; ****<0.0001.

### Study approval

All methods were carried out in accordance with relevant guidelines and regulations. All experimental protocols were approved by the respective licensing committees. Skin biopsies were obtained from healthy human volunteers under ethical approval of the Committee of Ethics and Academic and Scientific Deontology, Ministry of Education and Scientific Research, University of Medicine and Pharmacy of Craiova, Romania (Number 117/27.05.2015). Skin irritation experiments were performed in accordance with the guidelines set out by LaGeSo, project number EA1/1855/17. Informed consent was obtained from all the subjects participating to the study.

Mice experiments were performed in accordance with guidelines set out by LaGeSo, project number Reg 0222/16.

Zebrafish and embryos were raised and maintained according to standard protocols [71]. Experiments at the MPIIB were approved by, and conducted in accordance with, the guidelines set out by the State Agency for Health and Social Affairs (LaGeSo, Berlin, Germany). The Vivarium at NMS|FCM-UNL is licensed for animal work by DGAV, complying with the European Directive 2010/63/UE and the Portuguese Decree Law Number 113/2013, following the FELASA guidelines and recommendations concerning laboratory animal welfare.

## Acknowledgments

The authors thank Mary Louise Grossman and Souraya Sibaei for excellent editorial assistance, and Philippe Saikali for technical help (Department of Immunology, Max Planck Institute for Infection Biology, Charitéplatz 1, D-10117 Berlin, Germany). We also thank Sabrina Hadam, Eva Katharina Barbosa Pfannes and Annika Vogt for fruitful discussions (Department of Dermatology and Allergy, Clinical Research Center for Hair and Skin Science, Charité-Universitätsmedizin Berlin, Germany). We acknowledge Anne Diehl (Leibniz-Forschungsinstitut fuer Molekulare Pharmakologie, Berlin, Germany) for mice liver protein preparation. The zebrafish mpeg.mCherryCAAX SH378 mpx:GFP i114 line was kindly provided by Dr. Stephen Renshaw, The University of Sheffield, UK (http://www.sheffield.ac.uk/iicd/profiles/renshaw).

## Author contributions

L.L. and P.M.A. designed the study and performed the majority of experiments; T.D., J.W., H.J.M. and M.B. performed microarray analyses; A.K. and G.K performed *in silico* analysis. J.F. performed binding studies. C.C. and A.J. performed and analyzed zebrafish wound healing experiments; I.S., S.B.U. and I.M performed human skin biopsies experiments; R.H., M.K., U.G.B., U.Z., A.B.K. and M.S. performed experiments; M.L.M. supervised mouse experiments; F.S. and M.M. supervised the mouse wound healing experiments and performed contact skin irritation experiments; S.H.E.K. proposed and supervised project; L.L., P.M.A. analyzed data and wrote manuscript with major input from S.H.E.K.

## Data avalaibility

All data generated or analyzed during this study are included in this published article (and its Supplementary Information files). If additional details are desired, they are available from the corresponding author on request.

